# Real-time image processing toolbox for all-optical closed-loop control of neuronal activities

**DOI:** 10.1101/2020.06.22.166140

**Authors:** Weihao Sheng, Xueyang Zhao, Xinrui Huang, Yang Yang

**Affiliations:** School of Life Science and Technology, ShanghaiTech University, Shanghai 201210, China

## Abstract

The development of *in vivo* imaging and optogenetic tools makes it possible to control neural circuit activities in an all-optical, closed-loop manner [1], but such applications are limited by the lack of software for online analysis of neuronal imaging data. We developed an analysis software ORCA (Online Real-time activity and offline Cross-session Analysis), which performs image registration, neuron segmentation, and activity extraction at over 100 frames per second, fast enough to support real-time detection and readout of neural activity. Our active neuron detection algorithm is purely statistical, achieving much higher speed than previous methods. We demonstrated closed-loop control of neurons that were identified on the fly, without prior recording or image processing. ORCA also includes a cross-session alignment module that efficiently tracks neurons across multiple sessions. In summary, ORCA is a powerful toolbox for fast imaging data analysis, and provides a solution for all-optical closed-loop control of neuronal activity.

## Introduction

Establishing a causal relationship between neural activity and behavior is central to understand brain function, and this endeavor has been greatly facilitated by optogenetics [2–8]. Currently, most optogenetic experiments use pre-determined stimulation protocols, without considering the ongoing activity of relevant neurons. However, given that neuronal activities are highly heterogeneous—being closely linked to brain states and behaviors—it is clear that employing real-time activity-dependent stimulation protocols will better reveal the dynamic interactions between neural circuits and behavior [9–13]. Such “closed-loop experiments”, in which the input to the system (i.e., optogenetic stimulation) depends on the output (i.e., neural activity), are now within reach owing to optical imaging techniques for monitoring ongoing neural activities [14, 15]. However, accurately identifying active neurons in real time is an open challenge.

In priciple, identifying active neurons from raw imaging data requires two essential steps. The first is image registration, to compensate for the shift between image frames, for which several algorithms are available [16–18]; in this paper, we further accelerated this process by optimization. The second step and the major hurdle is to identify active neurons from the registered movies of ongoing activity. The current methods that feature either dimensionality reduction or deep learning are such that they require pre-processing of imaging datasets spanning a relatively long time for effective neuronal identification [19–24], and are therefore unsuitable for this purpose.

In addition, these imaging analysis pipelines cannot automatically track the same regions of interest (ROIs) across multiple sessions. Imaging the same population of neurons *in vivo* for an extended period of time is now possible with genetically encoded indicators and the implantation of chronic imaging windows or GRIN lenses [11, 25–27], but manually identifying the same neurons from multiple imaging sessions is time-consuming and error-prone. Additionally, some microscopes allow the user to rotate the optical axis. This feature adds flexibility to *in vivo* imaging, but at the same time exacerbates the difficulty of identifying the same neurons captured with slightly different angles, as rigid transformation alone cannot correct such distortions [23, 28].

We envisioned that using a statistical method based on calculating the temporal deviation of each pixel would allow us to achieve online identification of active neurons from streaming images. Pursuing this, we developed an image processing toolbox ORCA (Online Real-time activity and offline Cross-session Analysis), for fast image registration and online active neuron identification. ORCA also contains a cross-session alignment module for automatic tracking of neurons in long-term imaging (Figure 1). We demonstrate that ORCA can perform online identification of active neurons for closed-loop control, by identifying sound-responsive neurons in the mouse auditory cortex and selectively suppressing them with two-photon targeted optogenetic inhibition.

**Figure 1:**
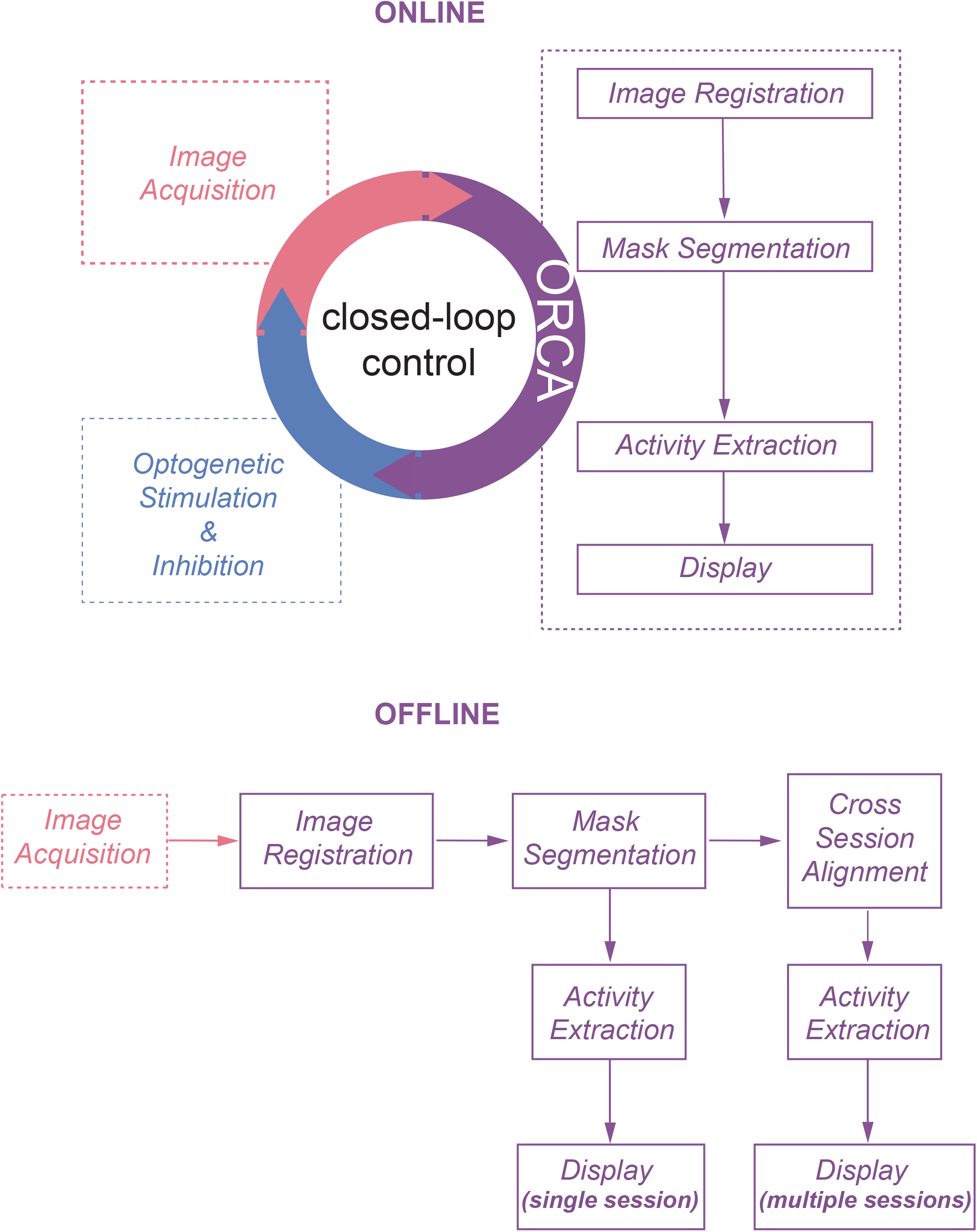
The online and offline pipelines of ORCA.

## Results

### Fast and accurate GPU-based image registration

The first module, *Image Registration*, removes motion artifacts caused by brain pulsation and body movement. To achieve high processing speed, we implemented an algorithm to search for the optimal translation optimized for GPU (Graphics Processing Unit) acceleration (see Methods for details). We compared the processing speed of ORCA with single-step DFT [17], TurboReg [16], and moco [18], using an example movie of 5040 frames of 512×512 pixels, containing both high-frequency lateral shifts and gradual drifting in the X-Y plane (Figure 2a). ORCA outperforms the other methods, especially when running on GPU (Figure 2b), with comparable registration accuracy as shown by the magnified z-stack images (Figure 2c, top panels). Examination of the offset of each frame relative to the template image indicated that ORCA captures both small and fast shifts, and large and slow drifts (Figure 2c, bottom panels).

**Figure 2:**
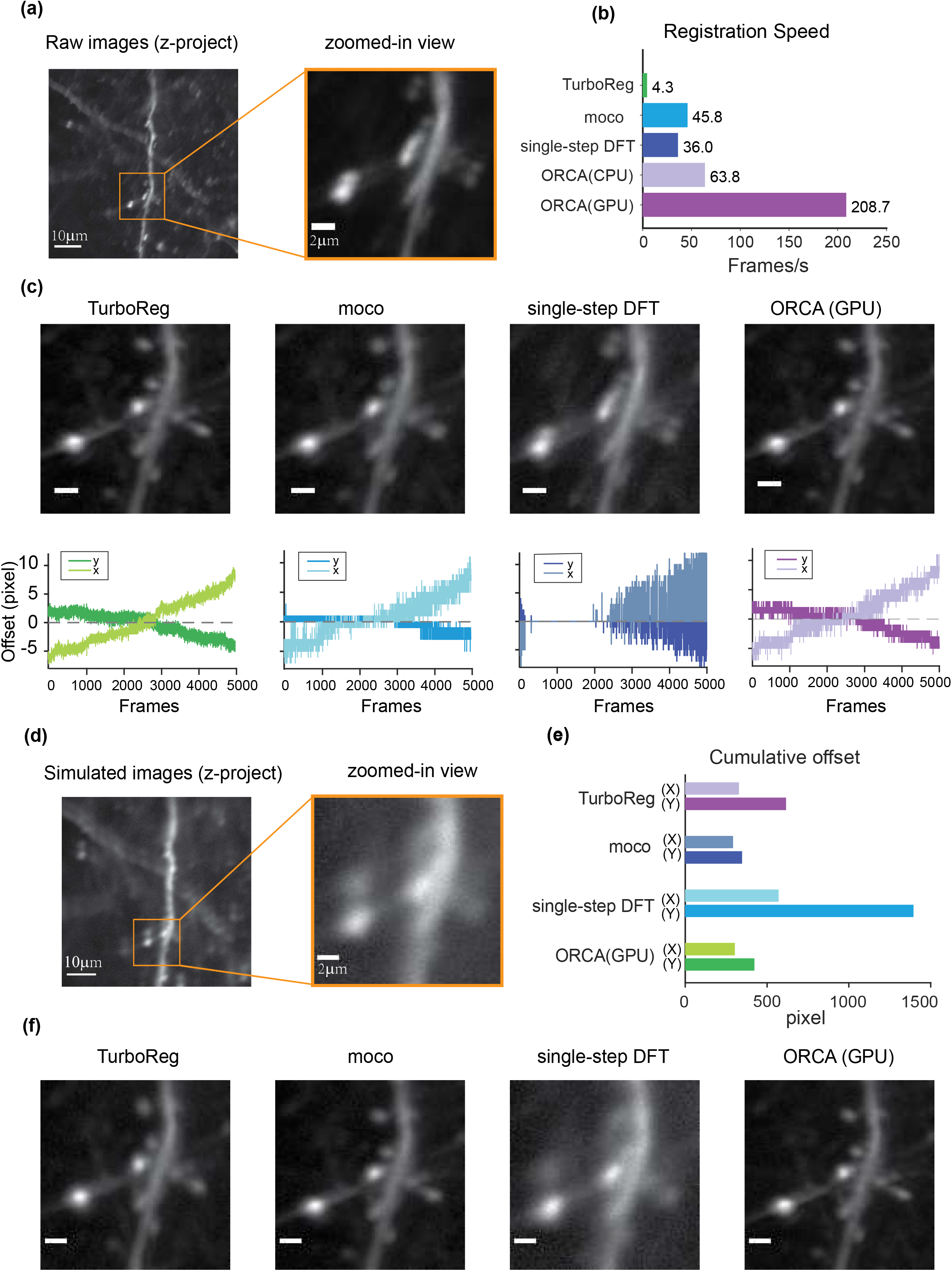
Raw image registration by ORCA and other softwares. (a) Z-project of raw images and zoom-in view. (b) Registration speed of different methods measured in frames per second. (c) Top panel: z-project of zoomed-in view after registration; bottom: calculated shifts in x & y direction. Scale bar, 2μm. (d) Z-project of simulated movie. (e) Quantification of registration accuracy with simulated data in (d). (f) Z-project of zoomed-in view after registration of simulated movie. Scale bar, 2μm.

In order to quantitatively assess the registration accuracy of ORCA, a simulated movie was created by compiling a series of identical images with pseudo-random x and y shifts, termed “true shift values” (Figure 2d). Image registration on this movie was conducted using ORCA and the other three methods above. ORCA achieves similar or better performance, as quantified by the discrepancy between calculated shifts and the ground truth, suggesting that ORCA is a more efficient tool in image registration (Figure 2e & f).

### Online identification and segmentation of active neurons

Most behavioral or neurophysiological experiments run in multiple trials. A typical trial lasts a few seconds, as does the inter-trial interval (ITI, Figure 3a). With a typical image acquisition rate of 10 ~ 30 frames/s, one trial generates a few dozen images. Adjusting optogenetic stimulation parameters based on neuronal activities in the preceding or the current trial requires identifying neurons and extracting their activities from a small set of images, and the computation must be completed within seconds. To satisfy such requirements, we developed a novel algorithm by summing temporal variance of each pixel and thresholding using Renyi’s entropy (Figure 3b, see Methods for details).

**Figure 3:**
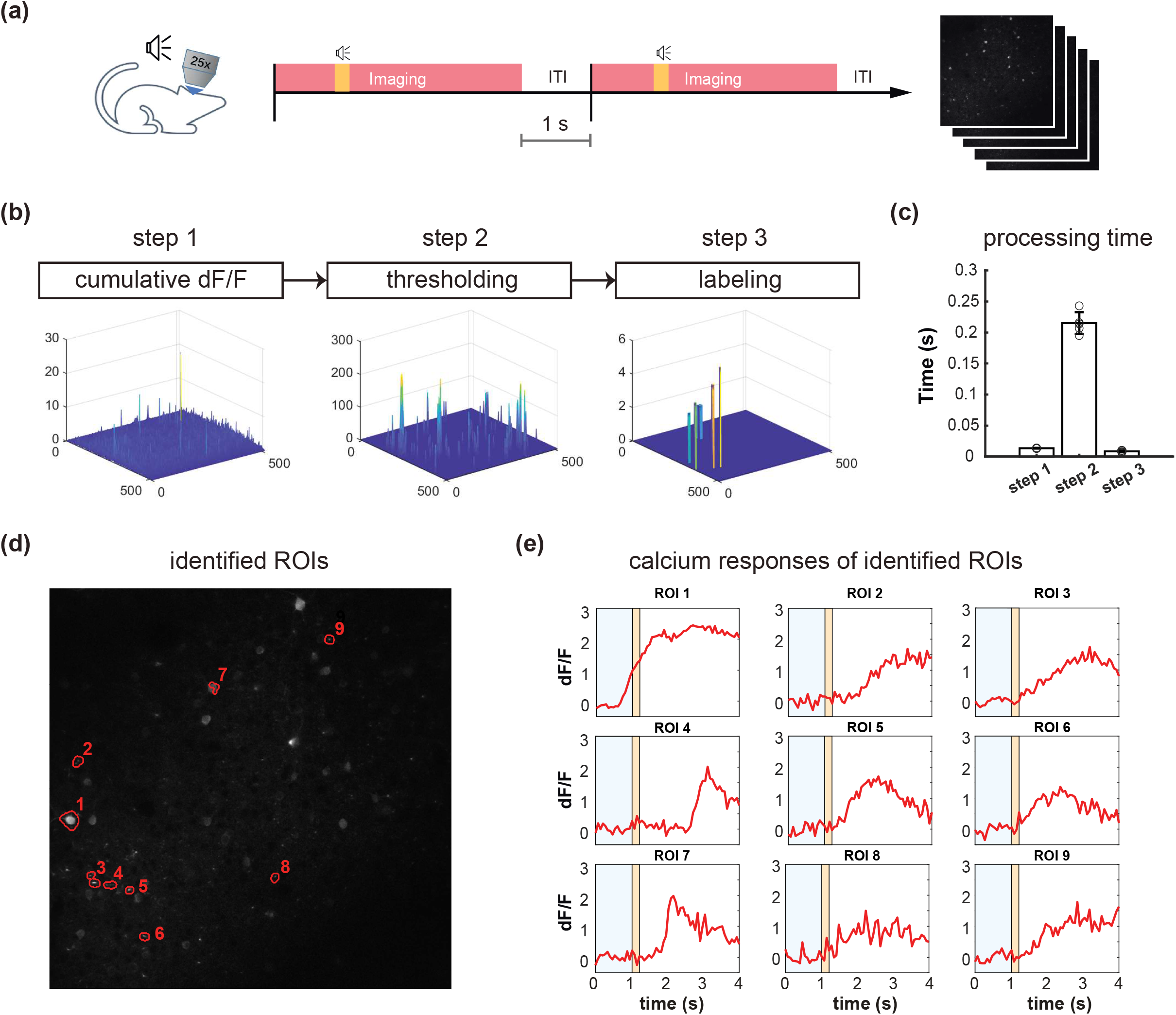
Online active neuron identification and activity extraction. (a) The flow of a typical trial-based imaging experiment. After 1-s baseline imaging, a pure tone lasting 0.2 s was played. Calclium activities were recorded for 4 seconds, and a 1-s inter-trial interval preceded the next trial. (b) Online neuron identification for each trial consists of 3 main steps. (c) Time spent in each step for processing data from one trial (60 frames). Each data point represents processing time for one trial. Average time is calculated for 5 trials. (d) An example of all identified ROIs in one trial. (e) Calcium responses (dF/F) of all ROIs identified in (d). Orange shading, pure tone.

To demonstrate online identification of active neurons, we performed *in vivo* two-photon calcium imaging in the mouse auditory cortex (ACx) and used ORCA to process the images. Each imaging trial lasted 4 s, including a 1-s “baseline” period and a 3-s “response” period. At an acquisition rate of 15 frames/s, each trial yielded 60 image frames. A 0.2-s tone was played after the baseline period (Figure 3a). As the mouse was anesthetized during imaging, shifting between frames was negligible during the 4-s imaging period, so we skipped *image registration* to further accelerate processing. The *Mask Segmentation* module first computed cumulative dF/F for each pixel in the response period, F defined as the average intensity of each pixel during the baseline period (Figure 3b, “cumulative dF/F”, see Methods for details). To separate major activities from background fluctuations, the second step was auto-thresholding of cumulative dF/F by Renyi’s Entropy [29] (Figure 3b, “thresholding”). To segment active neurons into regions-of-interest (ROIs), the third step was an additional thresholding based on user-defined ROI size and fluoresecent level (Figure 3b, “labelling”). The *activity extraction* module then extracted the activity trace of each identified ROI. The whole identification process took no more than 0.3 s (Figure 3c), well within the range of inter-trial intervals for sensory and behavioral experiments. Activity traces of all identified ROIs can be calculated and plotted instantly (Figure 3d & e), and can be exported to downstream control systems. We sorted the ROIs by their maximum dF/F in the descending order.

### Offline segmentation of active neurons for imaging sessions

For trial-based experiments, ORCA provides a fast offline image analysis solution. An imaging session with hundreds of trials can be analyzed within minutes using the algorithms for online identification, with one extra step to integrate ROIs identified from each trial to form a unified mask for the whole session (Figure 4a). For non-trial-based experiments, we incorporated a published algorithm, HNCcorr [20], for ROI segmentation. HNCcorr can also be used for trial-based experiments. We compared the processing speed and identification accuracy of ORCA and HNCcorr in an example trial-based imaging session. In this experiment, ACx neurons were imaged while a series of different pure tones were played. From this session, ORCA and HNCcorr identified 40 and 28 ROIs, respectively, with 17 ROIs identified by both (Figure 4c&d, Supp. Fig 1–2). ORCA identified more ROIs than HNCcorr, including some ROIs that showed significant calcium activity (dF/F) but were missed by HNCcorr (Figure 4c&d, ORCA ROI 13 and 15; Supp. Fig. 1). ROIs that were only identified by HNCcorr but were missed by ORCA did not seem to have significant calcium activity (Figure 4c&d, HNCcorr ROI 10 and 11; Supp. Fig. 2). Thus, ORCA is more effective in identifying active ROIs. Furthermore, the processing speed of ORCA is over 1000 times faster than HNCcorr (Figure 4b, 12 seconds compared to 4 hours, for 2880 image frames).

**Figure 4:**
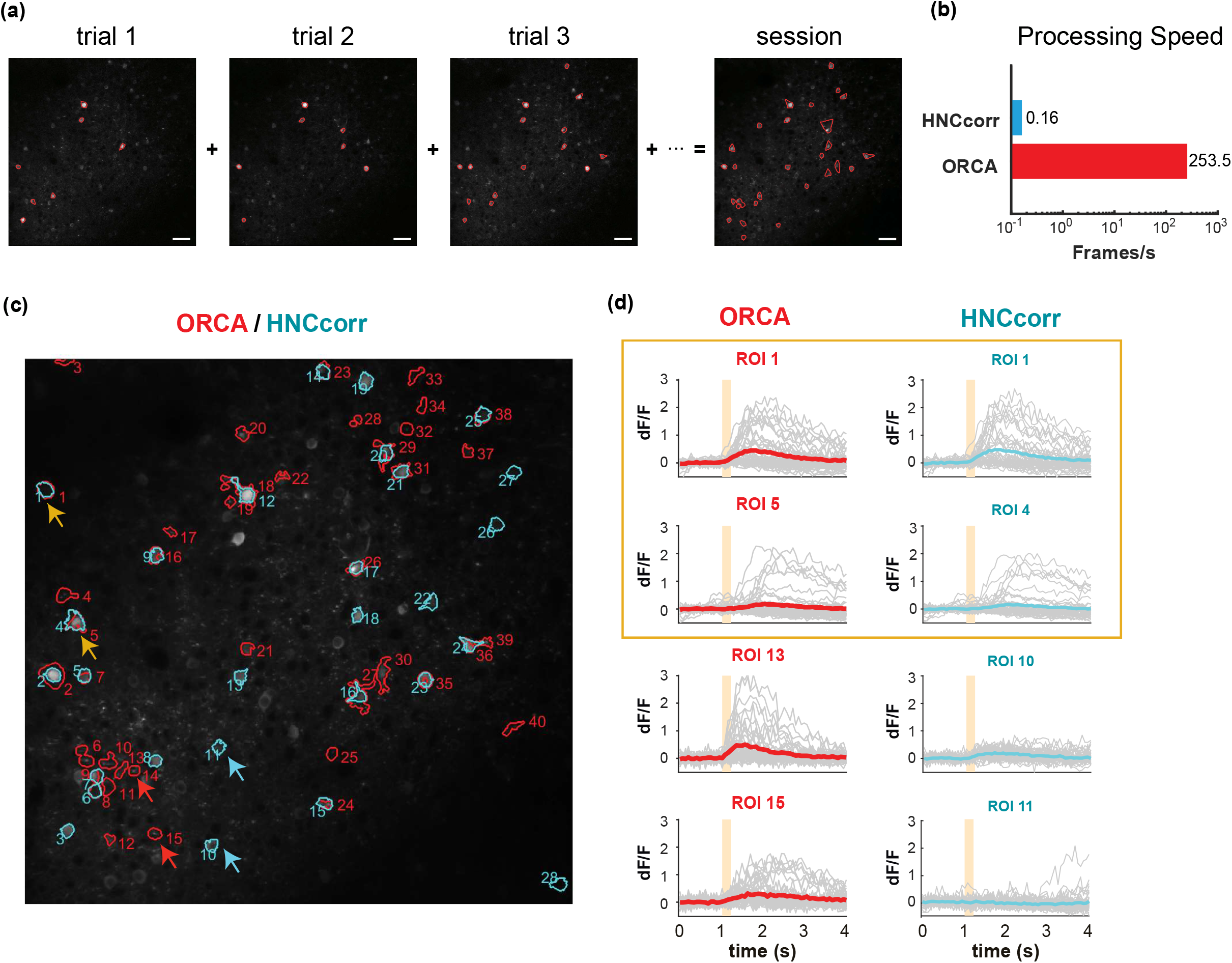
Offline active neuron identification and activity extraction for an entire session. (a) ROls were first identified in each trial, and then integrated to generate the session mask containing all active ROIs. (b) Processing speed of HNCcorr and ORCA for 512*512 image frames. (c) ROI segmentation results of the same imaging session by HNCcorr (blue) vs. ORCA (red). Orange arrows: examples of ROIs identified by HNCcorr and ORCA. Red arrows: examples of ROIs identified only by ORCA but missed by HNCcorr. Blue arrows: examples of ROIs only identified by HNCcorr but missed by ORCA. (d) Calcium activities of arrow-pointed ROIs in (c). ROIs 1 and 5 identified by ORCA corresponded to ROIs 1 and 4 by HNCcorr. ROIs 13 and 15 of ORCA were not identified by HNCcorr, and ROIs 10 and 11 of HNCcorr were not identified by ORCA.

### Cross-session alignment for repeated imaging

One challenge for long-term imaging is to track the same neurons across multiple imaging sessions obtained over an extended period of time (Figure 5a). Different sessions may have slightly different imaging angle due to imperfect adjustment of the microscope objective’s orientation relative to the plane of the imaging window, which hampers the identification of the same neurons across sessions. Furthermore, manual tracking is time-consuming and prone to human inconsistency. To address these issues, ORCA uses affine transformation, a linear mapping method that deals with uni-directional distortion, to correct for the angle differences between sessions (Methods). An example of two overlay imaging sessions of the same field of view (FOV) is shown in Figure 5b (left panel). The same neurons from different sessions may overlap completely or partially in the overlay image if they are in the center of the FOV, or not overlap if located in the periphery (Figure 5b, center panel). Our algorithm successfully aligned all the cells (Figure 5b, right panel). Masks from individual sessions were merged into one unified mask with a ‘capture-all’ strategy: all ROIs identified in any session were kept; for ROI identified in multiple sessions with similar (Figure 5c, blue arrow and box) or different shapes (Figure 5c, green arrow and box), the shape in the unified mask is the union of the shapes across all sessions. After indexing all ROIs in all imaging sessions, the *activity extraction* module extracts and plots the neural responses across different sessions. As an example, we plotted the individual and averaged changes in calcium signals (*ΔF/F*) of one ROI in two imaging sessions responding to pure tones of different frequencies (Figure 5d).

**Figure 5:**
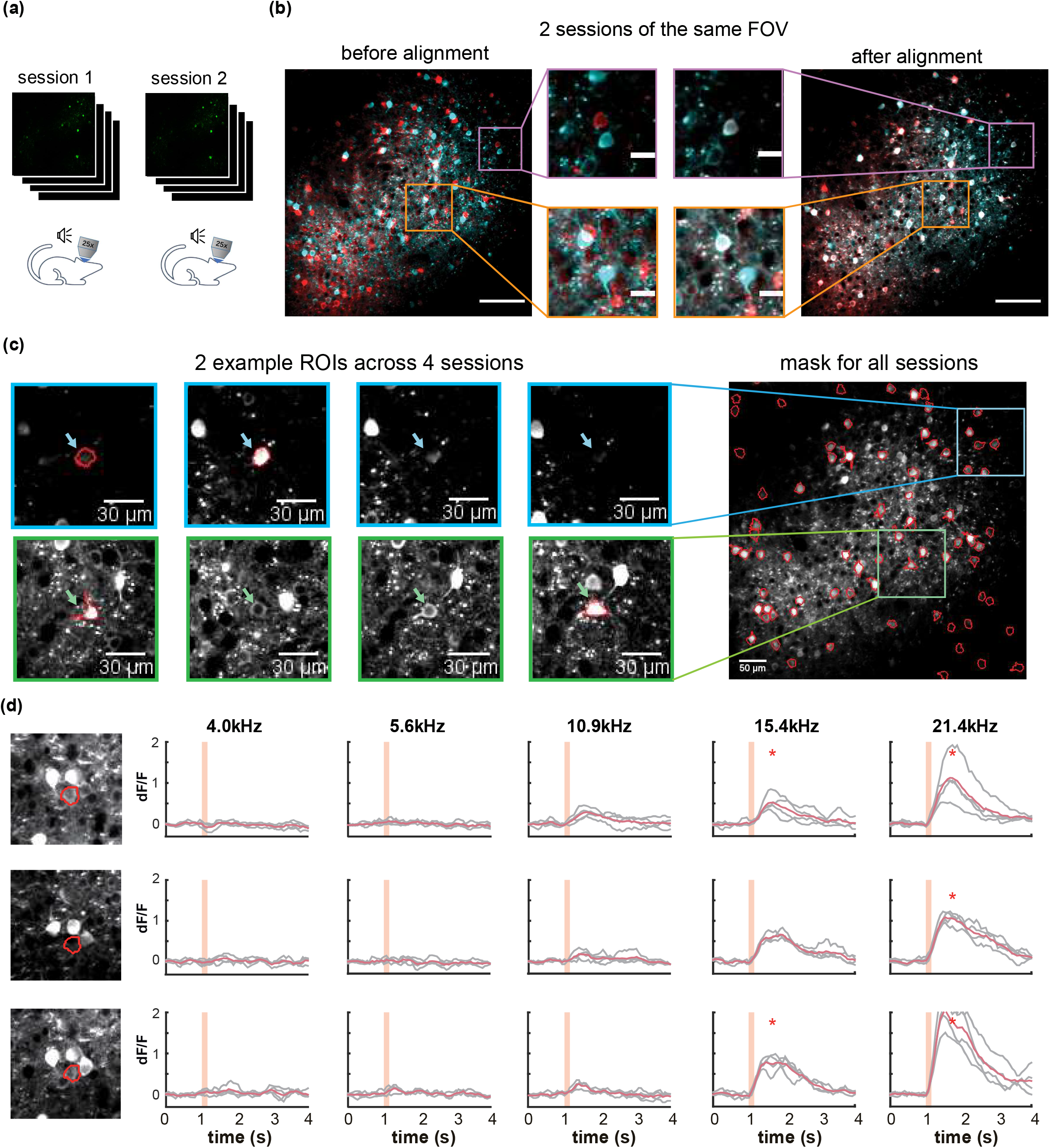
Multiple-session image alignment and data analysis. (a) Illustration showing multi-session imaging. (b) Cross session alignment for two sessions of the same FOV. Red and cyan colors are z-project images from two imaging sessions. Zoomed-in view on the left shows displacements of neurons at both center and margins, and on the right overlapping neurons after multi-session alignment. Scale bar, 100 μm for original image and 20 μm for zoom-in image. (c) Two example ROls were identified in 2 out of 4 sessions, with similar (blue) and different (green) shapes. They were included in the all-session mask. (d) Calcium responses of one example ROI across three sessions. Orange shading indicates pure tone duration.

### A demo for all-optical closed-loop control of neuronal activities

To demonstrate ORCA’s capacity for closed-loop photostimulation, we incorporated ORCA into a Thorlabs Bergamo II two-photon microscope with a spatial light modulator (SLM) for online manipulation of tone-responsive neurons in the mouse ACx (Figure 6a). We labeled ACx neurons with genetically encoded calcium indicator and optogenetic silencer by co-injecting AAV2/9-Syn-Cre, AAV2/9-hSyn-FLEX-GCaMP6s, and AAV2/9-hSyn-DIO-hGtACR1-mCherry viruses. A subset of neurons were co-labeled with GCaMP6s and GtACR1 (Supp. Fig. 3). To avoid photo-stimulation of GtACR1 while performing two-photon calcium imaging, we chose a wavelength (880nm) that could discriminate calcium-bound GCaMP6s from calcium-free GCaMP6s, and did not excite GtACR1 ([30–32], Methods).

**Figure 6:**
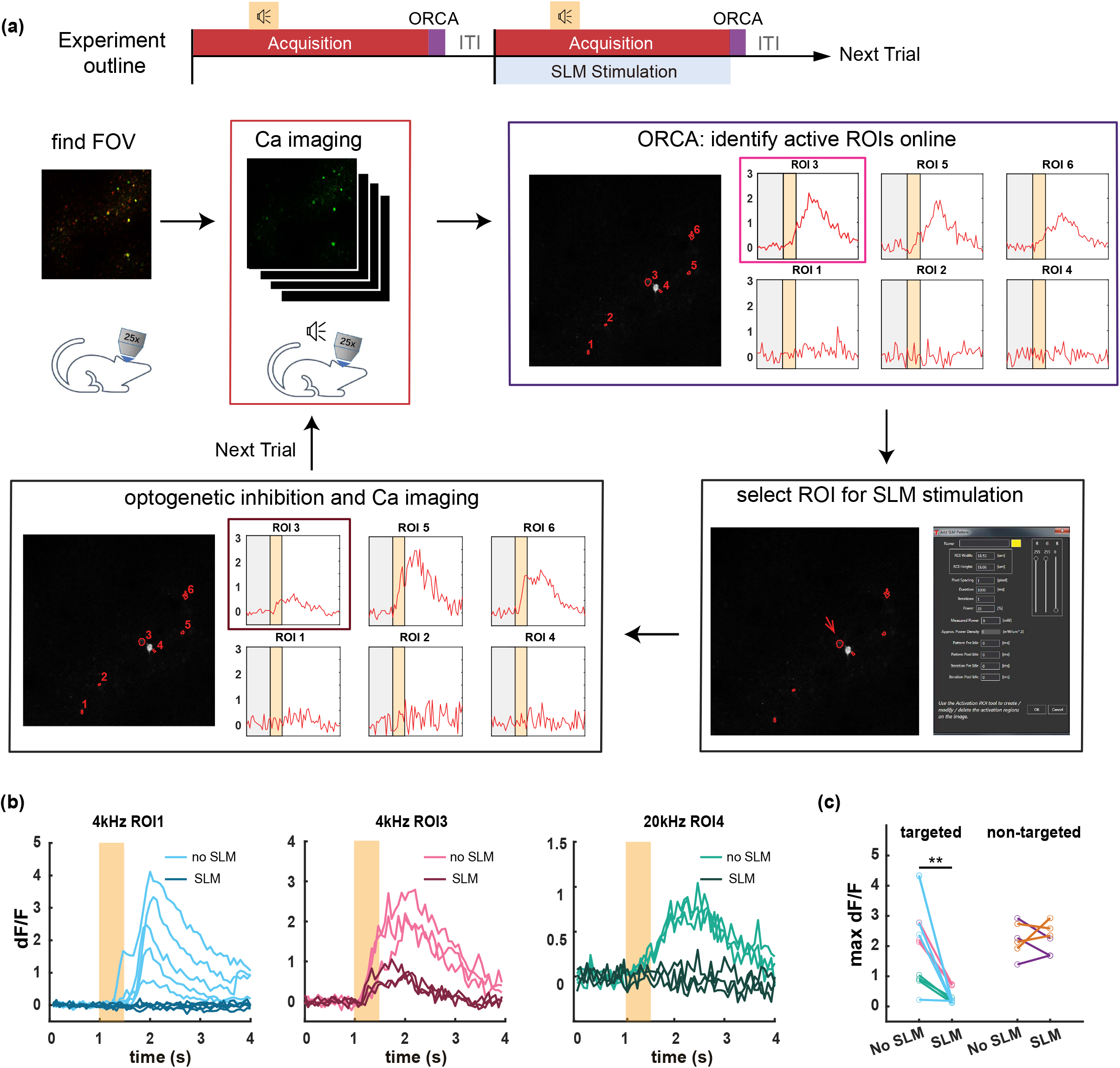
Application of ORCA in activity-based closed-loop experiments. (a) Timeline of the experiment. Two-photon calcium imaging is performed while different tones are played. Active ROIs are identified by ORCA online, and can be selected for SLM optogenetic inhibition. (b) Calcium responses of three different ROIs to 4kHz or 20kHz pure tones in SLM optogenetic inhibition trials (SLM) and non-inhibition trials (no SLM). (c) Maximum dF/F for ROIs selected for optogenetic inhibition (targeted), and ROIs that were not selected (non-targeted) in optically stimulated (SLM) and unstimulated (no SLM) trials.

We delivered a variety of pure tones (0.5 s each), and imaged neuronal calcium responses starting 1 s before the tone onset for a total of 4 s per tone presentation. The ITI was 1 s, common for trial-based sensory and behavioral experiments. We used ORCA to identify responsive ROIs and plot calcium responses online, before the onset of the next trial (Figure 6a). By loading identified ROIs into the stimulation software (ThorImage 4.3) before each stimulation trial, we demonstrated successful targeted optogenetic inhibition using SLM (Figure 6b&c, Supp. Figure 6) in stimulated but not control trials. The identification was proven accurate as in the stimulated trials, only the targeted ROIs were effectively inhibited, leaving the activities of the untargeted ROIs intact (Figure 6c). Thus, we showed that ORCA is well-adapted to commercial imaging and stimulation systems for activity-based closed-loop control.

## Discussion

Combining serveral novel algorithms, we developed ORCA, an imaging analysis toolbox that can process a small set of imaging data quickly for closed-loop neuronal stimulation, and perform cross-session image analysis for large datasets with high speed and accuracy. We validated our toolbox with calcium imaging experiments, but the software is in principle equally suitable for imaging studies using other fluorescent sensors, such as acetylcholine, dopamine and serotonin sensors [33–36].

To date, most optogenetic experiments are still open-loop, using stimulation parameters that are pre-determined rather than activity-guided. Only recently have neuronal activities been considered in guiding optogenetic manipulations, in the form of large-scale imaging data [12] or real-time activities of pre-identified neurons [13]. Nonetheless, an imaging analysis pipeline that identifies active neurons from ongoing imaging experiments, essential for effective manipulation of highly dynamic and heterogeneous neuronal populations, is still lacking. Such a pipeline needs to complete raw image registration, active component identification and activity extraction within a few seconds. Our GPU-based registration algorithm processes at >200 frames/s (512 ×512 pixels/frame). To our knowledge, only one method using an external OpenCV package written in C++ achieves comparable speed [24]. For online active component identification, our method is distinguished from previous methods [19, 21, 23] in that those methods require recording neuronal activity over an extended period and pre processing prior to online segmentation, whereas our method can identify active neurons on the fly at >200 frames/s. This feature improves the flexibility of neuron selection, which is especially valuable for experiments involving one-trial learning such as fear conditioning and novelty detection. Currently, we perform the second step in the *Mask Segmentation* module by calling an external function, the MIJ plugin of Fiji [37], which slows down the processing speed (Figure 3 b&c, *step 2*). In the future, by incorporating a Matlab function that calculates Renyi’s Entropy, we can further speed up this process.

Many studies have used long-term *in vivo* imaging to analyze the dynamics of the same population of neurons in multiple sessions. In such studies, neurons are typically tracked manually, which is error-prone especially with imaging angle differences. ORCA is the first to use spatial features of the original FOV to align different sessions with possible imaging angle discrepancies. Angle difference up to 5° can be corrected. Notably, angle differences that are too large will result in data loss especially in the margin, which cannot be fixed by post-processing. With its fast processing speed, the affline alighment function of ORCA can also be used for quality check and calibration during image acquisition. Users can first take a small number of images, calculate the angle offset from previous sessions, and adjust the microscope accordingly to match previous recordings. As more and more microscopes feature flexible imaging angles, ORCA is a good complement and widely applicable to these systems.

Segmenting neurons from noisy *in vivo* imaging data is challenging. To achieve high accuracy, segmentation methods often have to sacrifice processing speed and rely on large quantities of imaging data. In all-optical closed-loop experiments, however, data analysis requires fast processing using only small datasets. Using novel statistical algorithms, ORCA bridges the gap, and can be flexibly implemented into existing imaging systems to facilitate application of closed-loop manipulations owing to its modular design.

## Supplementary Figure legends

**Supplementary Figure 1.**
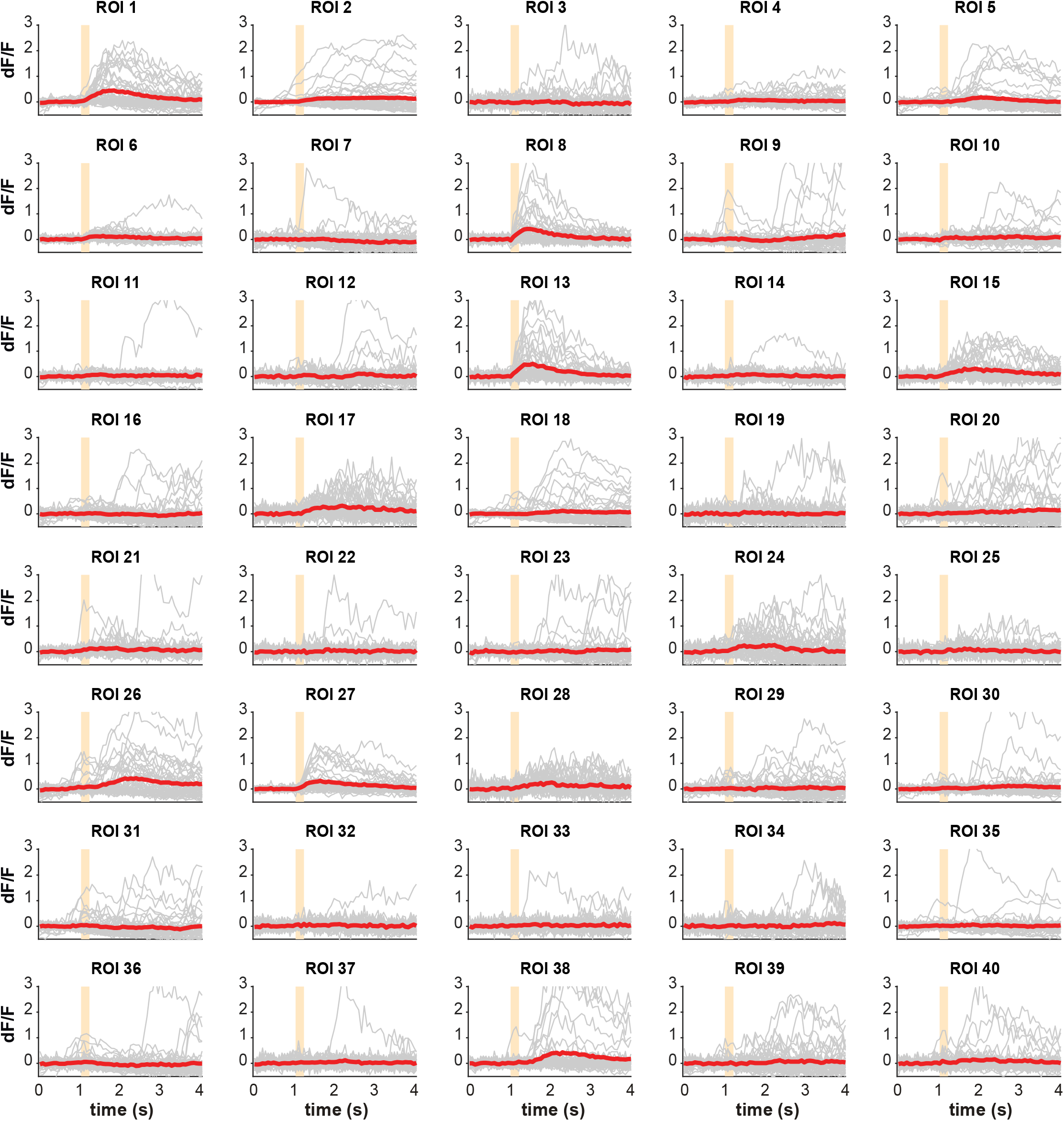
Calcium responses of each ROI identified by ORCA shown in Figure 4c.

**Supplementary Figure 2.**
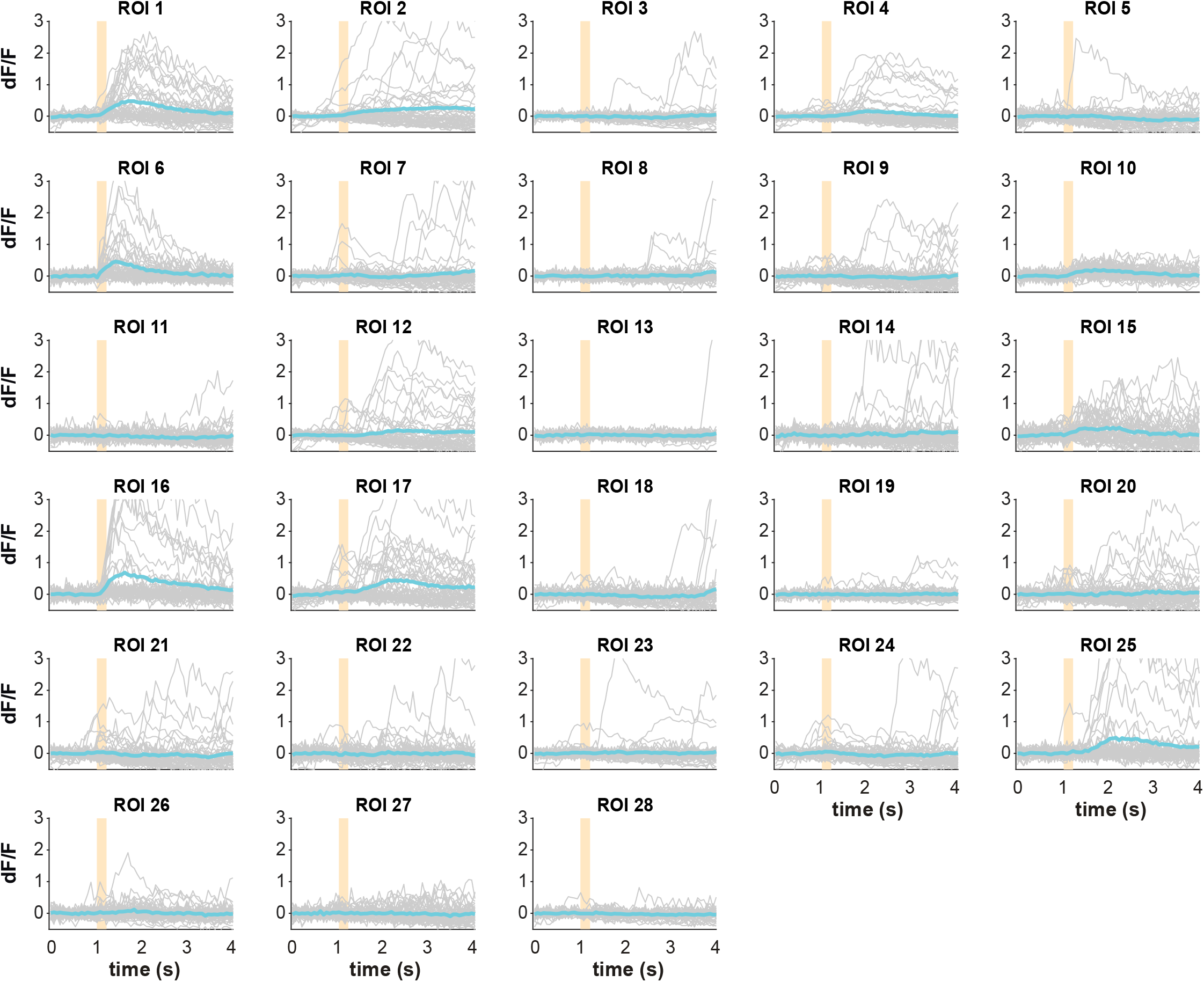
Calcium responses of each ROI identified by HNCcorr shown in Figure 4c.

**Supplementary Figure 3.**
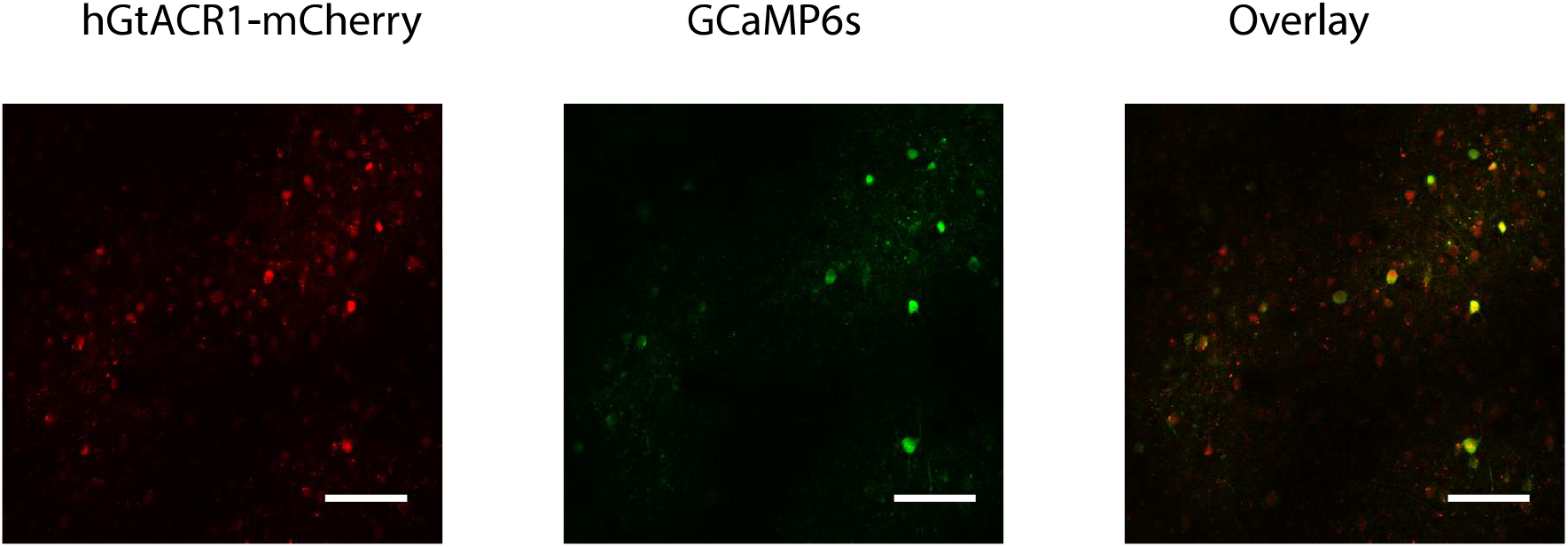
Expression of hGtACR1-mCherry (red), and GCaMP6s (green) in the FOV shown in Figure 6. Scale bar, 50 μm.

## Materials and methods

### Data availability

Code used in this paper is available at https://github.com/YangYangLab/ORCA.

### Animals

C57BL/6 mice were purchased from Slac Laboratory Animals (Shanghai, China). Mice were housed and bred in a 12 h light-dark cycle (7 am - 7 pm light) in the animal facility of ShanghaiTech University. Both male and female mice were used for the experiments. All procedures were approved by the Animal Committee of ShanghaiTech University.

### Virus injection

AAV2/9-Syn-Cre, AAV2/9-hSyn-Flpo-WPRE-pA, AAV2/9-hEFla-fDIO-GCaMP6s, AAV2/9-hSyn-FLEX-GCaMP6s, and AAV2/9-hSyn-DIO-hGtACR1-P2A-mCherry-WPRE-pA were purchased from Taitool Co., Shanghai, China. For virus injection, mice were anaesthetized with isofluorane (induction, 4%; maintenance, 1-2%) and positioned onto a stereotaxic frame (Reward Co., Shanghai, China). Body temperature was maintained at 37°C using a heat pad. Viruses were injected using a glass micropipette with a tip diameter of 15-20 μm through a small skull opening (< 0.5 mm^2^) with a micro-injector (Nanoject3, Drummond Scientific Co., Broomall, USA). Stereotaxic coordinates for auditory cortex (ACx): 2.46 mm posterior to the Bregma, 4.5 mm lateral from the midline, and 1.2 mm vertical from the cortical surface. For calcium imaging experiments, we mixed AAV2/9-hSyn-Flpo-WPRE-pA and AAV2/9-hEFla-fDIO-GCaMP6s, with the final titer of 1.3×10^12^ and 2.1×10^12^ viral particles per ml, respectively. For closed-loop stimulation experiments, we mixed AAV2/9-Syn-Cre, AAV2/9-hSyn-FLEX-GCaMP6f-WPRE-pA, and AAV2/9-hSyn-DIO-hGtACR1-P2A-mCherry-WPRE-pA, with the final titer of 6.4×10^9^, 6.9×10^12^ and 1.4×10^12^ viral particles per ml, respectively. We injected 0.2 μl virus mixture into the auditory cortex for all experiments, and waited 3-4 weeks before two-photon imaging experiments.

### Cranial window implantation

Mice were anaesthetized with isoflurane (induction, 4%; maintenance, 1-2%) and positioned onto a stereotaxic frame (Reward Co.). Body temperature was maintained at 37°C using a heat pad. Lidocaine was administered subcutaneously. The muscle covering the auditory cortex was carefully removed with a scalpel. A 2×2 mm^2^ piece of skull over ACx was removed, exposing the dura. The cranial window was sealed with a custom-made double-layered cover glass. UV-cure glue and dental acrylic were used to cement the cover glass onto the skull. A custom-made stainless steel headplate with a screw hole was embedded into the dental acrylic for head-fixed two-photon imaging.

### Two-photon calcium imaging and optogenetic stimulation using spatial light modulator

Mice were injected with pentobarbital sodium (20 mg/kg) and head-fixed using the implanted headplate. Image series were taken at 15Hz with a two-photon microscope (Bergamo II, Thorlabs, Newton, NJ, USA) equipped with a 25X/NA 1.05 objective (Olympus, Kyoto, Japan) and a Ti:sapphire laser (DeepSee, Spectra-Physics, Santa Clara, USA) tuned to 880 nm. Mice were imaged when a series of pure tones (4kHz, 12kHz, 20kHz) were played. Sound was controlled using a Bpod state machine (Sanworks Co., Rochester, USA) and open-source Bpod software (Sanworks Co.).

A fixed-wavelength laser (1040nm, Spectra-Physics, USA) and spatial light modulator (Thorlabs) were used for two-photon targeted optogenetic silencing. A cycle of 40 was used for inhibiting selected ROIs (0.1s per cycle).The SLM was controlled by ThorImage (Thorlabs) and the control module of ORCA.

### Hardware for imaging data analysis

We used a Dell workstation to perform all computations. The workstation is equipped with two Intel Xeon Gold 5122 CPU, 64GB DDR4 RAM, and an Nvidia Quadro P6000 GPU, running Linux Debian 10. ORCA uses GPU to accelerate most processing steps, while CPU versions are also provided in the package. We recommend the GPU version for best performance.

### Preparation and evaluation of image registration

To compare the speed of different algorithms, we performed registration in their default environments: single-step DFT and ORCA were tested in MATLAB R2020a (MathWorks, Natick, USA), while TurboReg and moco in Fiji (ImageJ 1.48v, Java 1.6.0_24, 64bit). File read/write time was excluded for better evaluation of computational efficiency of each algorithm. Timing of TurboReg, single-step DFT and ORCA started after files were loaded into the memory, and stopped as soon as algorithms produced registered results. Timing of moco was evaluated using a screen recording software (Kazam Screencaster) and inspected manually.

To quantitatively compare different algorithms, we generated a simulated movie as follows. An example movie consisting of 1200 frames was repeatedly registered using moco, TurboReg (accurate mode) and ORCA for 10 times each. This movie was then carefully examined by tracking major features in the images and measuring their movements frame-by-frame, to ensure it reached maximum possible stability. Then, periodical shifts caused by heartbeats were simulated by adding directional movements using the following equation:

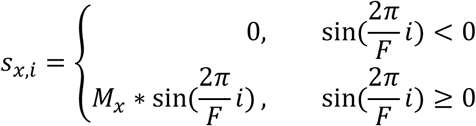

where *F* indicated frames-per-second during acquisition, *M_x_* the maximum shifts in x direction, and *i* the current frame count. This kept all positive values in *s_x_* and set all negative offsets to zero. An additional random distortion was also introduced using pseudo-random numbers as offsets. Thus shifts in x direction, *s_x_*, became

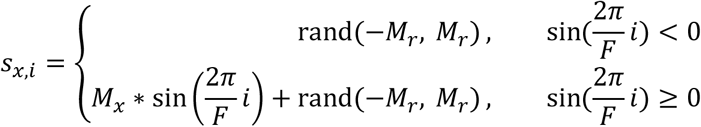

where rand(−*M_r_*, *M_r_*) generated a pseudo-random sequence ranging from [−*M_r_*, *M_r_*]. True shift values were generated in both x and y direction in the same manner with varying parametres, and simulated movie was tested among all four registration methods.

Registration accuracy was quantified as

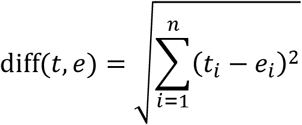

where *t* was the true values, and *e* was the estimated shifts generated by each algorithm. The registration accuracy was calculated separately for x and y directions.

### Image registration algorithms in ORCA

Inspired by the moco algorithm [18], we combined multiple computational ideas to produce a faster implementation. Let *α* be an image frame of *h* rows (height) and *w* columns (width), and α_i,j_ denote the pixel of the *i*-th row, *j*-th column. Let *t* be the template image (usually the first frame in an image series). Images *t* and *α* are standardized by mean-centering before registration:

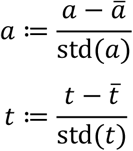

Let *Ms* be the user-defined maximum shift value. We set its default value to 1/5 of the frame size. Users are advised to start with the default setting and adjust the parameter from there, as appropriate choice of *Ms* is important for the best performance. An *Ms* too large (e.g., for a 512*512 pixel image, *Ms* = 170) may result in faulty matches, while an *Ms* too small (e.g., for a 512*512 image, *Ms* = 1) will cause the registration to fail, because the x-y shift between frames is likely to exceed 1 pixel.

We search for the optimal *x,y* offsets within the maximum shifts to minimize the difference between *α* and *Tm* in their overlapping region, defined as

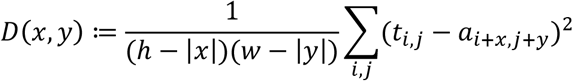

Substituting the denomiator (*h* – |*x*|)(*w* – |*y*|) with *A_x,y_*, the area of overlap between the template and the target frame, we can rewrite the equation as:

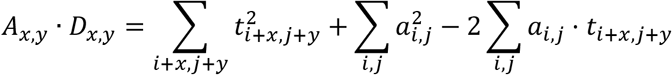

in which *D_x,y_* is the sum of all pixels squared in the overlapping area of the template, and *i,j* are corresponding indices. Given that 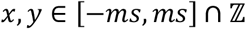, there are finite possibilities of these values, and they can be calculated for any given *x,y* in the range beforehand.

To efficiently calculate these values, we register template *t* onto current frame *a*, so that ∑*t*^2^ only need to be calculated once throughout the whole registration process. ∑*a*^2^ and ∑*α*·*t*, are both related to the current frame *α*. Notice that, once we determined the maximum shift *ms*, only a central fraction of the original images are then registered to the template. To further accelerate the computation, we crop out the marginal areas of the original image, using only the central part to register against the full template. The cropped image, *α*’, should be

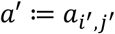

Where *i*′ ∈ [*Ms, h – Ms*], and *j*′ ∈ [*Ms, w – Ms*]. Thus we rewrite the equation into:

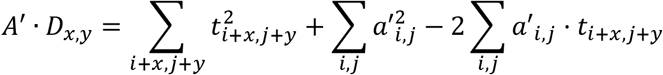

In this equation, *α*’ are the same cropped image for any valid 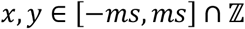, and the overlapping area *A*’ becomes static. The only dynamic part in the equation thus becomes ∑*α*·*t*. ∑*α·t* is the convolution of *a* and 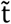 evaluated at (0,0). Letting 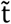 be the rotation of *t* by 180 degrees, we have

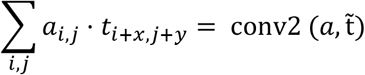

Convolution can be efficiently calculated using two-dimensional fast Fourier transform (2D-FFT):

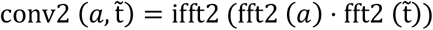

and thus we developed a highly optimised conv2 function (conv2_`fftvalid.m` in the source code) for our registration, which is much faster than MATLAB’s default implementation using basic addition and multiplication.

∑*α*^2^ can also be calculated as

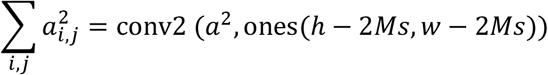

where ones(*h – 2Ms, w – 2Ms*)) is a matrix of size (*h – 2Ms, w – 2Ms*) with all elements being 1, in the MATLAB notation.

The whole registration process can be further accelerated by downsampling and doing a local search after upsampling. Running on GPU, our code is much faster than most other registration algorithms.

### Online trial-by-trial identification of active neurons

We combined several statistical tools for online segmentation of active ROIs on a trial-by-trial basis, each trial consisting of *N* frames of size (*H*, *W*) that constitute an image stack.

The dF/F (here denoted by 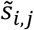 for pixel (*i,j*)) is computed by subtracting the baseline intensity 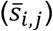 of each pixel in the image stack and divided by 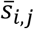:

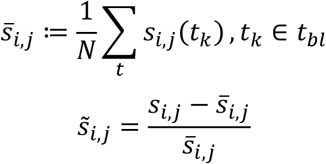

where *s_ij_*(*t*) represents a pixel (*i*, *j*) of the *t*^th^ image in the stack, *t_bl_* is the baseline period (defined by user). Then, dF/F of each pixel is summed up to form a two dimensional time-average image. Values below zero are substituted by zeros.

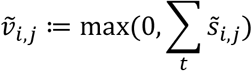

Here, 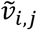 represents the cumulative dF/F for pixel (*i*, *j*) in the 2-D image. To increase the signal-to-noise ratio, we use standard deviation of 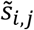 as a scaling factor, and multiply by 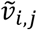:

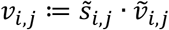

The product matrix *v_i, j_* undergoes a filter that converts it to a binary image using Renyi Entropy-based auto thresholding [29] by calling the MIJ plugin in Fiji [37]. The binary image is smoothed using a Gaussian filter to combine neighbouring pixels. Two additional user-defined thresholds can be applied to determine whether to keep the identified ROIs (area threshold, default set at 16 pixels; intensity threshold, default set to 10). Pixels containing 1 (“positive pixels”) indicates a masked region.

### Offline manual segmentation of components in a movie

Since automated identification may falsely identify unwanted components or neglect true active neurons, we provide a user-friendly interface for manual labelling of active neurons in a given movie, incorporated in the mask_segmentation module. Our manual labelling program allows users to play the movie for full length or just one trial, and add or erase components.

### Cross session alignment

For multi-session imaging data of one FOV, we use time-average images to represent the imaging sessions, and define the time-average image of the first imaging session as the reference image. In a three-dimensional space, we define the xy plane as the plane of the reference image, and z axis perpendicular to xy. We then define the target image to be the time-average of a different imaging session. For sessions without rotation on the z axis, displacements in x and y axis were corrected simply by moving target images in x, y direction for optimum alignment. For sessions with rotation, because the image plane preserves the collinearity of the FOV, displacements can be corrected using affine transformation. We first computed similarity between the reference image and the target image, then applied different transformations onto the target image and search for maximum similarity between them. We used Matlab function `imregtform()` with ‘`rigid`’ or ‘`affine`’ parameters, and applied the transformation matrix by the function. We incorporated manual inspection and refinement by the users in this module. After user confirmation of the alignment results, masks with segmented ROIs from different sessions were combine to generate a superset of ROIs. These ROIs were then transformed to the original session based on respective transformation matrices. ROIs in the margins may be cropped out in the cross-session alignment.

### Activity extraction

As ORCA extracts neuronal activity trial-by-trial, the user needs to supply information about the trial structure, in particular timing parameters. The baseline activity *F_0_* of each ROI is the average of all pixels over a user-defined time window before stimulus onset:

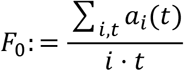

where *α_i_* is a pixel in the corresponding ROI, and *t* ∈ [Baseline Frames]. Then, the normalized activity of all ROIs is calculated by subtracting and dividing baseline activities:

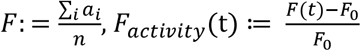

## Acknowledgement

We thank Drs. J. Lu, M. Ho, S.T. Han and M. Tsakiris for critical reading and editing of the manuscript. We thank the imaging facility of School of Life Science and Technology, ShanghaiTech University, for technical support. We also thank the following funding agencies (2018YFC1005004 and NSFC31970960 to YY) for the support of the work.

